# RSM01, an extended half-life RSV monoclonal antibody with a high barrier to resistance

**DOI:** 10.64898/2026.02.04.703721

**Authors:** Jonne Terstappen, Emanuele C. Gustani Buss, Jason S. McLellan, Vicky-Lynne Baillie, Louis J. Bont, Mauricio T. Caballero, Klasina Chappin, Janissa J.A. van Duijn, Robert Jan Lebbink, Isabelle C.E. van Lotringen, Joris J.T. Roggekamp, Anouk Versnel, Marco C. Viveen, Hanneke J.A.A. van Zoggel, Eveline M. Delemarre, Aurelio Bonavia, Natalie I. Mazur

## Abstract

Respiratory syncytial virus (RSV) causes substantial infant morbidity and mortality, particularly in low- and middle-income countries (LMICs). RSM01 is a long-acting RSV monoclonal antibody (mAb) in development for LMICs. To date, RSM01 resistant mutants have not been able to be generated *in vitro*. We systematically assessed the *in vitro* resistance barrier of RSM01 relative to licensed RSV mAbs and the susceptibility of a panel of global contemporary strains to RSM01. The emergence of mAb-resistant mutant (MARM) was assessed for RSM01, nirsevimab, and palivizumab. Triple plaque-purified RSV-A2 and RSV-B1 were serially passaged under mAb pressure to generate MARMs, which were phenotyped and genotyped. Moderate resistance was defined as > 3-fold and high resistance > 30-fold increase in IC_50_ compared with parental strains. The susceptibility of contemporary clinical strains from South Africa, Argentina, and the Netherlands to RSM01 was tested in a neutralisation assay. RSV-A and B lab strains developed high resistance to palivizumab and nirsevimab, while RSM01 pressure induced moderate resistance. No MARMs demonstrated a fitness advantage; one RSM01 MARM incurred fitness costs. RSM01 potently neutralized global contemporary RSV-A and B strains. In conclusion, *in vitro* RSM01 resistance was infrequent and moderate, suggesting a high resistance barrier *in vitro*. Laboratory-selected MARMs may not directly predict clinical escape, but the confirmation of the high barrier to resistance of RSM01 is encouraging and provides an alternative in the event that resistance is observed with widespread use of current licensed anti-RSV mAbs.

**One Sentence Summary:** *In vitro* RSV resistance to RSM01 is infrequent and moderate, suggesting a high barrier to resistance.

## INTRODUCTION

Respiratory syncytial virus (RSV) is the most common cause of lower respiratory tract infections (LRTI) among infants and young children and a major contributor to global child mortality (*1, 2*). While new immunisation products have the potential to substantially improve infant health, access to monoclonal antibody (mAbs) remains highly inequitable (*3, 4*). The mAb nirsevimab (*5*) has been implemented almost exclusively in high-income countries (HICs) (*6*), despite more than 97% of the RSV-associated infant mortality occurring in low- and middle-income countries (LMICs) (*1*). Ensuring a durable, affordable, and resistance-resilient RSV mAb is therefore essential to achieving global health equity.

Current and developmental mAbs neutralise RSV by binding the fusion (F) protein and blocking viral entry into host cells. RSM01 is a fully human IgG1 mAb with half-life extending YTE-mutation targeting antigenic site Ø on the prefusion (preF) conformation of the RSV F protein (*7*). It demonstrates potent neutralization of diverse RSV-A and RSV-B isolates in vitro, prophylactic efficacy in animal models, a favourable non-clinical pharmacology profile, and was well tolerated in healthy adults volunteers in a phase 1 clinical study (*8*). The development program explicitly prioritizes manufacturing efficiency and affordability, positioning RSM01 as a potential single-dose, season-long RSV prevention strategy tailored for LMIC settings.

RSM01 and nirsevimab bind overlapping epitopes within antigenic site Ø (*8, 9*), whereas clesrovimab targets site IV on both pre- and post-fusion state of the F protein (Fig. S1) (*10*). Together, these mAbs are replacing palivizumab, which targets site II and has a restricted indication and requires multiple costly injections per season (*11*). As the global use of long-acting mAbs increases, understanding their vulnerability to viral escape is critical. Like other RNA viruses, RSV can generate monoclonal antibody-resistant mutants (MARMs) under selective mAb pressure, as seen in *in vitro* and animal studies (*8, 12-18*). Resistance-associated substitutions are rarely observed in circulating strains (∼1%) (*17, 19-21*) and often impose viral fitness costs (*14, 16, 17*). To date, *in vivo* resistance in infants immunised with palivizumab or nirsevimab remains uncommon (*22, 23*) and breakthrough infections are often linked to suboptimal dosing (*24, 25*). Although some resistance-associated mutations have been reported, mainly in RSV-B (*23, 25*), circulating viruses have largely remained susceptible (*23*), and target epitopes are highly conserved (*18, 21*). The long-term effectiveness of antibody-based prevention depends on the barrier to resistance of each product, however, because nirsevimab has only recently been introduced, the clinical significance of resistance cannot yet be fully assessed.

High-barrier-to-resistance mAbs (those requiring multiple or fitness-impairing mutations for escape) are particularly important in settings where genomic surveillance is limited and timely product replacement is challenging and sometimes economically inviable. Evaluating resistance robustness during development is therefore a critical component of sustainable RSV immunoprophylaxis. Previous studies failed to generate escape mutants against RSM01 or its parental antibody, while MARMs were readily obtained for palivizumab and nirsevimab (*8*); however, more stringent experimental design and quality control are required to validate these findings.

Here, we systematically assess the resistance barrier of the LMIC-focused candidate RSM01 relative to existing RSV mAbs by characterizing *in vitro*–selected escape mutants, their effects on viral fitness, the prevalence of associated mutations in circulating RSV sequences. We also present the susceptibility of newly isolated contemporary global strains to RSM01. Together, these data provide an integrated assessment of the resistance potential of RSM01 and its suitability for durable global RSV prevention.

## RESULTS

### Selection and characterisation of RSV MARMs

#### *In vitro* generation of MARMs

In order to understand potential immune escape, we generated MARMs for RSV-A2 and B1. ATCC master stocks were triple plaque purified (TPP) into parental strains (Fig. S2). ATCC RSV-A2 master stock contained a minor variant N262Y mutation in comparison with the master stock consensus sequence (allele frequency (AF) = 31%) in the palivizumab epitope, while 36% of the RSV-A2 parental strain contained the mutation (Table 1). ATCC RSV-B1 master stock contained a K65E mutation, which was lost during plaque purification. TPP parental strains showed comparable growth curves to the ATCC master stocks (Fig. S3). Parental strains were serially passaged under mAb pressure for 10 passages under various *in vitro* culture conditions (Fig. 1A-B and S2). We display experimental conditions previously tested to generate *in vitro* MARMs against RSM01 (Fig. 1C) alongside the experimental conditions tested in this study to show a full overview of the potential of RSM01 to generate MARMs *in vitro* (*8*).

**Table 1.** Genotypic and neutralisation changes in selected MARMs.

| Virus name | Subtype | mAb pressure<br>(concentration in ng/mL) | Passage | Amino acid substitution |  | Mean IC<sub>50</sub><br>(ng/mL) | Fold-change to<br>parental strain |
| --- | --- | --- | --- | --- | --- | --- | --- |
|  |  |  |  | Consensus strain | Minor variant (AF) |  |  |
| Virus stocks |  |  |  |  |  |  |  |
| A2 ATCC Master stock | RSV-A2 | None | ATCC+2 | None | N262Y (31%) | .. | .. |
| A2 TPP Parental | RSV-A2 | None | ATCC+7 | None | N262Y (36%) | RSM01 20.3<br>Nirsevimab 31.3<br>Palivizumab 3460.8 | 1 |
| A2 virus controls<sup>a</sup> | RSV-A2 | None | Parental+10 | N262Y | None | .. | .. |
| B1 ATCC Master stock | RSV-B1 | None | ATCC+1 | K65E | None | .. | .. |
| B1 TPP Parental | RSV-B1 | None | ATCC+9 | None | None | RSM01 42.8<br>Nirsevimab 101.2<br>Palivizumab 2116.0 | 1 |
| B1 virus control | RSV-B1 | None | Parental+12 | None | None | .. | .. |
| RSM01 MARMs |  |  |  |  |  |  |  |
| RSM-1 | RSV-A2 | Fixed RSM01 (11.5) | Parental+12 | I206T/N262Y | None | 352.1 | 17.3 |
| RSM-2 | RSV-A2 | Rising RSM01 (46) | Parental+12 | I206T/N262Y | None | 220.9 | 10.9 |
| RSM-3 | RSV-B1 | Rising RSM01 (230) | Parental+15 | N216H<sup>b</sup> | None | 26.5 | 0.6 |
| Nirsevimab MARMs |  |  |  |  |  |  |  |
| NIR-1 | RSV-A2 | Fixed nirsevimab (30) | Parental+12 | D194N/N262Y | None | 120.4 | 3.8 |
| NIR-2 | RSV-A2 | Rising nirsevimab (96) | Parental+9 | N63K/N262Y | None | 154.5 | 4.9 |
| NIR-3 | RSV-B1 | Fixed nirsevimab (15) | Parental+12 | K65E | None | >124 500 | >1230.2 |
| NIR-4 | RSV-B1 | Fixed nirsevimab (30) | Parental+12 | K65E | N63D (17%) | >124 500 | >1230.2 |
| NIR-5 | RSV-B1 | Rising nirsevimab (1500) | Parental+12 | K65E | None | >124 500 | >1230.2 |
| RSM-4 | RSV-B1 | Fixed RSM01 (23) | Parental+12 | K65E | None | >124 500 | >1230.2 |
| Palivizumab MARMs |  |  |  |  |  |  |  |
| PAL-1 | RSV-A2 | Fixed palivizumab (50 000) | Parental+12 | N262Y | None | >2 500 000 | >722.4 |
| PAL-2 | RSV-B1 | Fixed palivizumab (2000) | Parental+13 | K272I | None | >2 500 000 | >1181.5 |

ATCC RSV-A2 and RSV-B1 master stocks were triple plaque purified to generate parental strains, which were serially passaged under mAb pressure. Mab concentrations were either fixed for 10 passages or increased based on CPE. Parental strains served as references for MARM sequence analysis; minor variants were reported at 5-40% allele frequency with $\geq 200\times$ read depth. Rising mAb represent the concentration at MARMs characterisation. <sup>a</sup>All four virus controls showed the same mutation. <sup>b</sup>No mutation was observed in NGS after viral propagation. An N216H substitution observed by Sanger sequencing in two passages (p8, p13) was absent in reverse reads and an intermediate passage (p10), indicating it was not stably fixed. AF, allele frequency; IC<sub>50</sub>, half-maximum inhibitory concentration; mAb, monoclonal antibody; MARMs, monoclonal antibody resistant mutants; RSV, respiratory syncytial virus; TPP, triple plaque purified; .., not available.

**Fig. 1.**
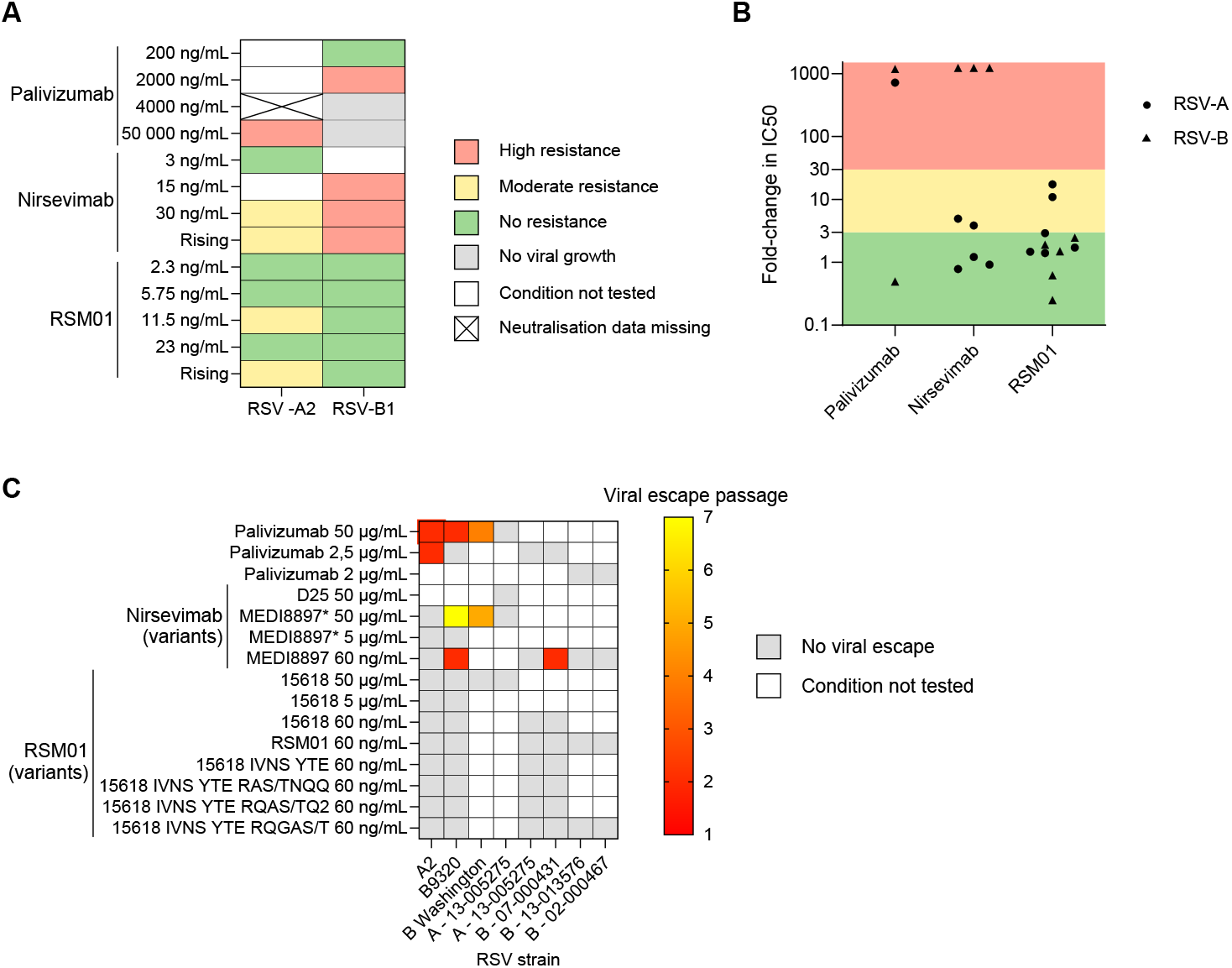
Generation of RSV mAb resistant mutants. **A.** RSV-A2 and B1 mAb resistant mutants (MARMs) were generated in the present study by 10 serial passages under mAb pressure. Mab concentrations (fixed or rising) are presented on the y-axis and RSV strains on x-axis. Neutralisation data for RSV-A2 cultured under 4000 ng/mL palivizumab is missing (cross). **B.** All potential MARMs with >3 fold increase in IC_50_ compared with parental strains as determined in a neutralisation assay were further neutralised according to principles of mass violation. Green, yellow, and red banding represent varying degrees of resistance (no, moderate, and high resistance, respectively) in terms of IC_50_ fold change compared with parental strains. Mean IC_50_ values from two independent assays are presented. **C**. Generation of RSV MARMs in an independent laboratory (Arsanis Biosciences GmbH) using various laboratory and clinical strains at various concentrations of mAb (*8*). Several variants of nirsevimab (MEDI8897) were used, where MEDI8897 represents nirsevimab without YTE mutation and D25 is the parental antibody. RSM01 variants included parent antibody 15618 IVNS YTE and other 15618 variants. Viral escape was defined as CPE as observed under light microscopy, confirmed by F protein sequencing. No viral escape (grey) was observed for RSM01 and variants, while viral escape was readily observed for both palivizumab and nirsevimab during serial passage (red-yellow scale, where red signifies early passages and yellow at later passages). CPE, cytopathic effects; IC_50_, half-maximum inhibitory concentration; mAb, monoclonal antibody; MARMs, monoclonal antibody resistant mutants; RSV, respiratory syncytial virus.

#### Identification and characterisation of MARMs

All virus strains cultured under mAb pressure at passage 10 as well as strains showing high cytopathic effects (CPE) under rising mAb pressure at earlier passages, were systematically screened for at least 3-fold increase in half inhibitory concentration (IC_50_) compared with parental strains. Selected MARMs were further characterised by neutralisation and sequencing to align functional with genotypical changes (Table 1 and Fig. 1A-B). Additionally, RSV-B1 grown under RSM01 pressure resulted in a nirsevimab MARM with a known mutation in the nirsevimab epitope (K65E) and was also further characterised. Virus controls, cultured without mAb pressure, showed a N262Y mutation as the consensus strain in four RSV-A2 controls at passage 10 and no mutations occurred in F protein for one RSV-B1 control at passage 12. MARMs were named after the mAb used to generate the MARM and numbered.

#### RSM01-MARMs

RSM01 pressure generated 3 MARMs out of 10 conditions (30% escape frequency; Fig 1B). Out of all RSM01 MARMs, 2/3 showed moderate resistance. RSM-1 and -2, RSV-A2 cultured under fixed 11.5 ng/mL RSM01 and under rising RSM01 concentration (up to 46 ng/mL, see Materials and Methods) (Table 1), showed the same mutations (I206T/N262Y) and moderate resistance (10.9 and 17.3-fold increase in IC_50_). Moreover, RSM-3 was observed as a MARM during culture as it showed >10% CPE under high RSM01 pressure (230 ng/mL). However, no genotypical or functional resistance was observed after viral propagation (0.6-fold change in IC_50_), potentially due to a loss in mutation during propagation or due to missed mutations outside the F-protein. Sanger sequencing of earlier passages did not reveal consistent consensus-level mutations across passages: an N216H mutation was detected in the forward read of two passages (passage 8 and 13) but was absent in the corresponding reverse reads and in intermediate passage 10, potentially due to an unstable mutation.

#### Nirsevimab-MARMs

Nirsevimab pressure induced resistance in 5/6 (83%) growth conditions and 3/5 MARMs (60%) showed high resistance and 2/5 (40%) showed moderate resistance (Fig. 1B-C and Table 1). Two RSV-A2 MARMs, NIR-1, with the D194N mutation neighbouring the nirsevimab epitope, and NIR-2 (N63K mutation) both showed moderate resistance (3.8 and 4.9-fold increase in IC_50_, respectively). NIR-3 to NIR-5, RSV-B1 cultured under three concentrations of nirsevimab, all showed the same K65E mutation in the nirsevimab binding site, which was associated with complete neutralisation escape (>1230 fold increase in IC_50_). One of these isolates (NIR-5) showed an additional minor variant at position N63D (AF=17%), with an uncertain effect on neutralisation as the K65E MARM showed complete escape. Notably, a MARM cultured under RSM01 pressure (RSM-4) with the same K65E mutation was still susceptible to RSM01 but showed escape against nirsevimab (>1230 fold increase in IC_50_).

#### Palivizumab-MARMs

Under palivizumab pressure, 2 out of 5 growth conditions (40%) resulted in escape; both MARMs showed high resistance (Fig. 1B-C and Table 1). The RSV-A2 MARM (PAL-1) cultured under fixed 50 µg/mL palivizumab showed a N262Y mutation in the consensus, which conferred complete neutralisation escape (>722-fold increase in IC_50_). RSV-B1 cultured under fixed 2000 ng/mL palivizumab showed a K272I mutation (PAL-2), also associated with complete palivizumab escape (>1181-fold increase in IC_50_).

#### MARMs growth kinetics

To determine the impact of antibody escape mutations on viral growth, we characterised the growth kinetics of TPP parental RSV-A2 or B1 and their derived MARMs by inoculating Hep-2 cells with an MOI of 0.01. The cultured supernatant was sampled twice daily for four days and viral growth was assessed using a TCID_50_ assay. Parental RSV-A2 and derived MARMs show similar growth kinetics and peak titres (Fig. 2A-C). RSM-3 had slower *in vitro* growth and lower viral titres compared to the parental strain (Fig. 2D). The B1 nirsevimab-MARMs showed similar viral growth to the parental strain (Fig. 2E). Viral growth kinetics of RSM-4 are missing as it was initially not identified as a resistant mutant to RSM01 but showed genotypic and phenotypic resistance to nirsevimab in cross-resistance testing. Overall, viral resistance conferred no advantage or fitness costs as the parental strains and derived RSM01 or nirsevimab-MARMs exhibited comparable or slower growth kinetics and similar peak titres.

**Fig. 2.**
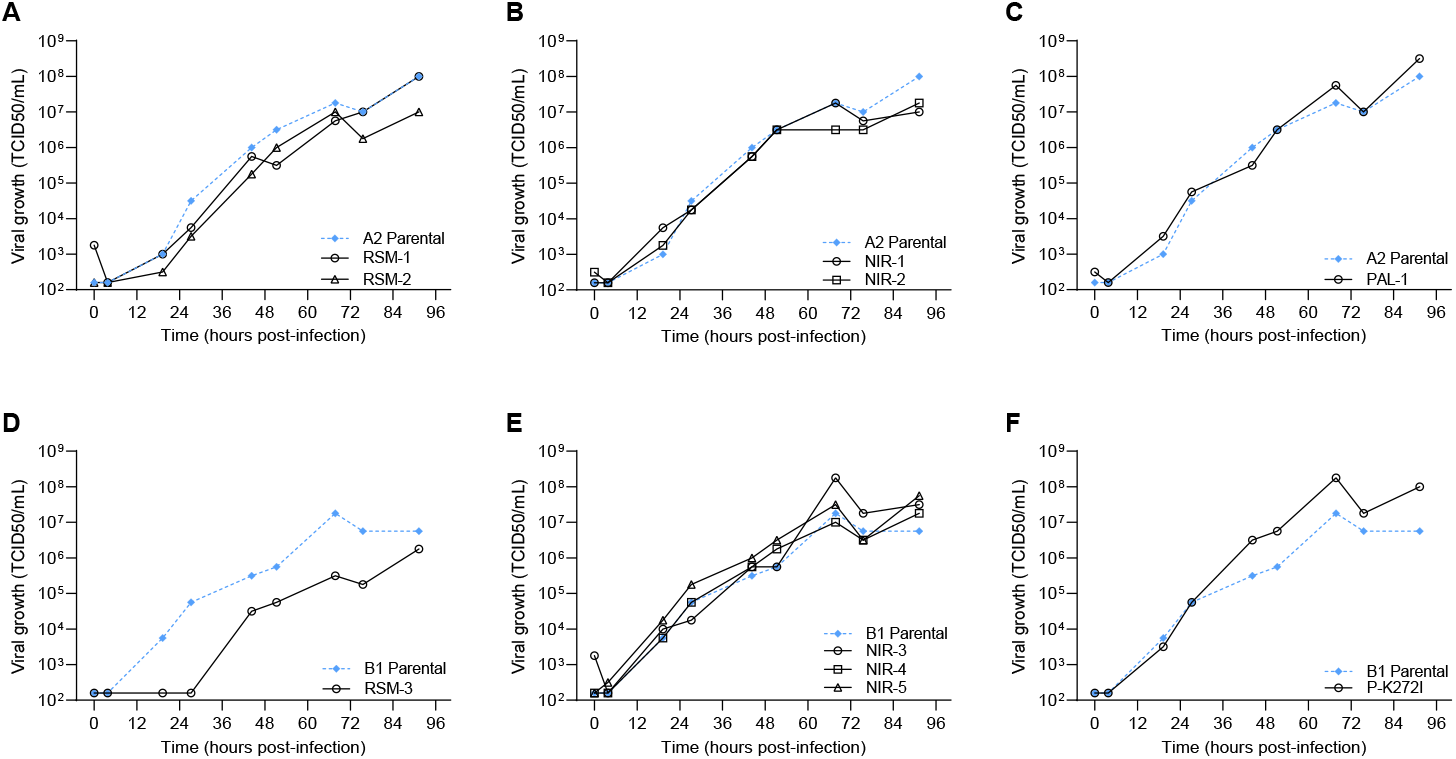
Viral growth kinetics of MARMs on HEp-2 cells. Growth kinetics of parental TPP RSV-A2 and B1 and the derived MARMs. HEp-2 cells were inoculated at an MOI of 0.01 TCID_50_/cell. The cultured supernatant was sampled twice daily for four days and viral growth was assessed using a TCID_50_ assay. RSV-A2 MARMs (**A-C**) and RSV-B1 MARMs (**D-F**) show similar growth to their parental TPP strains, with the exception of RSM-3 (**D**), which shows delayed and lower titre growth. MARMs, monoclonal antibody resistant mutants; MOI, multiplicity of infection; RSV, respiratory syncytial virus; TCID_50_, half-maximum tissue culture infective dose.

#### Cross resistance of MARMs to other RSV mAbs

All MARMs could be rescued by at least one other mAb. Three RSM01 MARMs (RSM-1 to RSM-3) could still be neutralised by nirsevimab and clesrovimab (Fig. 3). RSM-4 was not resistant to RSM01 or clesrovimab, but was highly resistant (>1230 fold-change) to nirsevimab despite culture under RSM01 pressure. All nirsevimab MARMs could be rescued by RSM01 and/or clesrovimab.

**Fig. 3.**
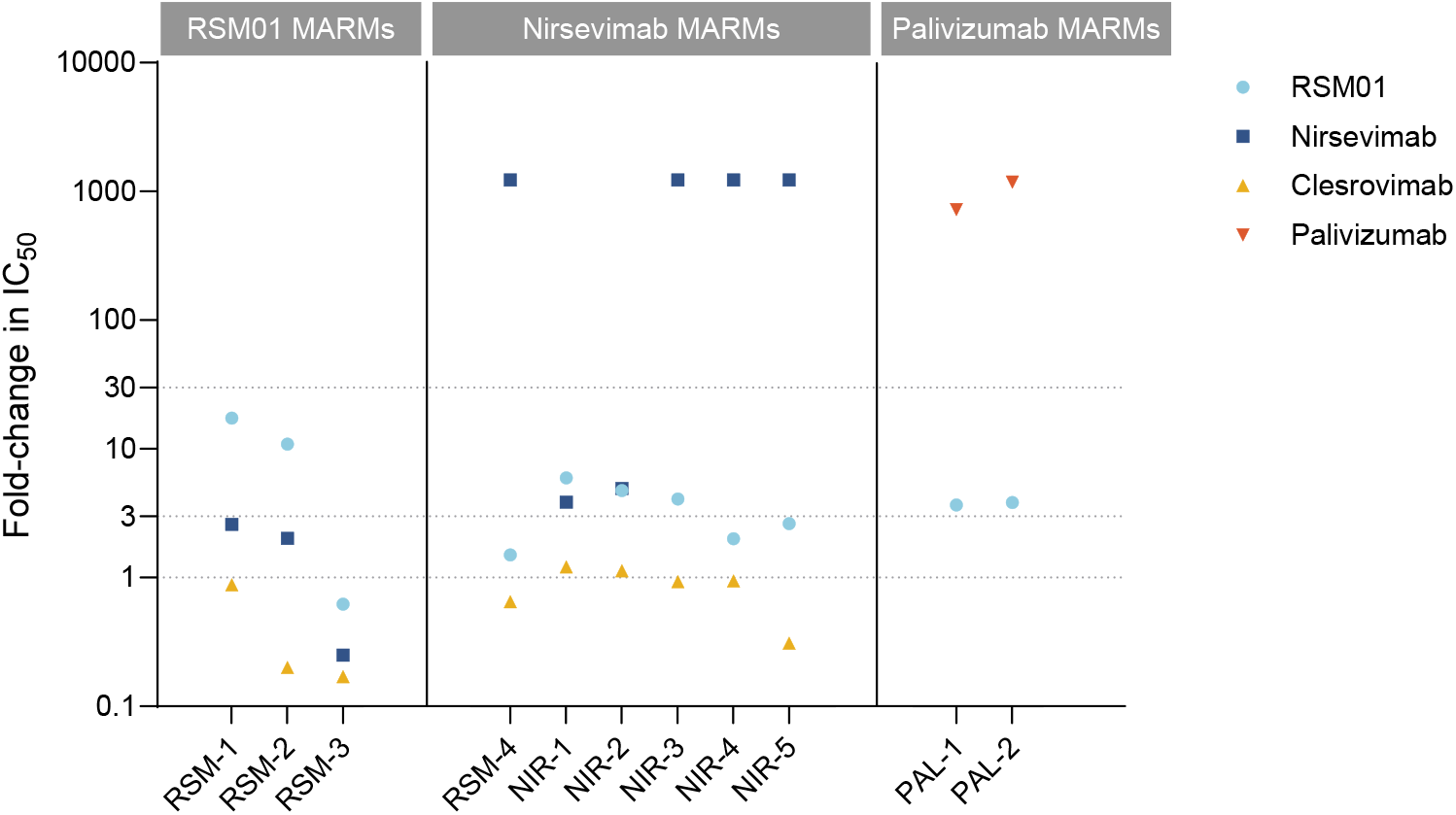
Cross-resistance of MARMs against other RSV mAbs. *In vitro* neutralisation of MARMs by other available RSV mAbs. Potency is expressed as fold-change in IC_50_ compared with the parental strain per mAb. IC_50_, half-maximum inhibitory concentration; mAb, monoclonal antibody; MARMs, monoclonal antibody resistant mutants; RSV, respiratory syncytial virus.

#### Prevalence of resistance-associated mutations in circulating strains

To investigate the occurrence of MARMs in currently circulating RSV strains, we compared the seven amino acid substitutions identified in the 11 *in vitro* MARMs to a global GISAID database (Table 2). All polymorphisms occurred at low frequencies (ranging between 0.04 and 0.16%) in current global strains. The I206T mutation associated with moderate resistance against RSM01 (and nirsevimab) occurred at a frequency of 0.16%. Notably, K272I, which arose in a palivizumab *in vitro* RSV-B MARM, was detected exclusively in RSV-A clinical strains.

**Table 2.** Natural occurrence of polymorphisms identified in MARMs.

| Mutation | RSV-A |  | RSV-B |  |
| --- | --- | --- | --- | --- |
|  | Sequences with mutation/Total | Frequency (%) | Sequences with mutation/Total | Frequency (%) |
| N63D | 0/5037 | 0 | 2/5120 | 0.04 |
| N63K | 0/5037 | 0 | 0/5120 | 0 |
| K65E | 0/5037 | 0 | 2/5120 | 0.04 |
| D194N | 0/5037 | 0 | 0/5120 | 0 |
| I206T | 8/5037 | 0.16 | 0/5120 | 0 |
| N216H | 0/5037 | 0 | 1/5120 | 0.02 |
| N262Y | 0/5037 | 0 | 0/5120 | 0 |
| K272I | 8/5037 | 0.16 | 0/5120 | 0 |

Whole genome sequences were extracted from GISAID database (May 5, 2023 to Dec 1, 2025). MARMs, monoclonal antibody resistant mutants; RSV, respiratory syncytial virus

### Amino acid variability in mAb binding sites by Shannon entropy analysis

Shannon entropy analysis was performed to assess diversity on the surface F glycoprotein. The F protein of the RSV-A isolates showed 27 sites with high entropy (H > 0.1) (Table S2), of which two are located within epitopes: the nirsevimab binding site at residue 63 (H=0.11) and the palivizumab binding site at residue 276 (H=0.59) (Fig. 4A). RSV-B isolates displayed 17 highly variable positions, two of which are located in overlapping RSM01/nirsevimab binding sites (Table S3; Fig. 4B).

**Fig. 4.**
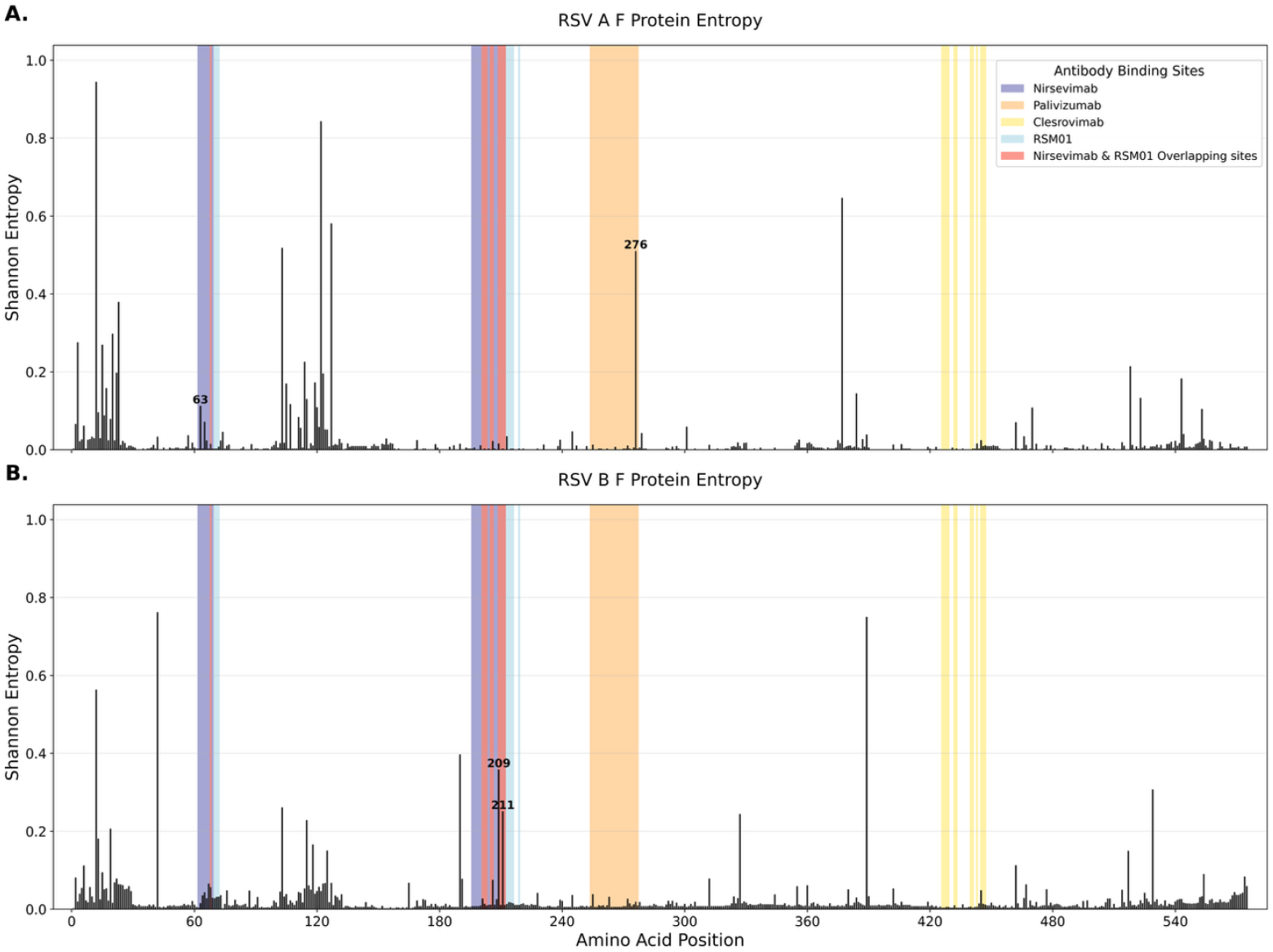
Shannon entropy across amino acids positions in RSV F-protein sequences. The RSV whole genomes were downloaded from GISAID (May 5, 2023, up to Dec 5, 2025) with 5037 RSV-A and 5120 RSV-B F gene sequences. Shannon entropy was estimated based on each amino-acid position across the F-protein and proportion of amino acids found at a given residue for all genomes for both RSV-A (**A**) and RSV-B (**B**). The F protein structure is color-coded to indicate mAb binding sites (nirsevimab in dark blue, RSM01 in light blue, overlap between nirsevimab and RSM01 in salmon, palivizumab in orange, and clesrovimab in yellow). High entropy residues (H≥0.1) in mAb binding sites are annotated.

The RSM01 binding site was largely conserved across both RSV subtypes, with the majority of positions showing H < 0.1 (Fig. 4). However, within the region where RSM01 and nirsevimab binding sites overlap, residues 209 (H=0.35) and 211 (H=0.25) exhibited elevated polymorphisms among circulating RSV-B strains (Fig. 4B). Additionally, in RSV-A the nirsevimab binding site showed increased entropy at residue 63 (H=0.11) and the palivizumab binding site displayed higher entropy at residue 276 (H=0.51) (Figure 4A).

### Susceptibility of contemporary global strains to RSM01

RSM01 was further evaluated for its neutralizing activity against a panel of contemporary strains in a neutralisation assay. In total, 31 out of 46 isolates (14 RSV-A and 17 RSV-B) were successfully cultured on HEp-2 cells with sufficiently high titres for the neutralisation assay (14 from South-Africa, 11 from Argentina, and 6 from the Netherlands).

All clinical isolates were characterised for subtype-specific diversity patterns. For RSV-A, seven distinct sublineages were identified: A.D.1.9 (n=5), A.D.3 (n=4), A.D.3.1 (n=1), A.D.3.3 (n=1), A.D.4.1 (n=1), A.D.5.1 (n=1), and A.D.5.2 (n=2) (Fig. 5A). RSV-B was characterised into three sublineages: B.D.E.1 (n=12), B.D.E.1.1 (n=4), and B.D.E.1.7 (n=2) (Fig. 5B). Phylogenetic trees were constructed from whole-genome sequences to assess the diversity of the panel for RSV-A and RSV-B contemporary strains from Argentina, South Africa, and the Netherlands, along with globally representative sequences from six geographic regions spanning 1950–2025 (Fig. S4). For both RSV-A and RSV-B, when compared with global context data, most study samples clustered with high statistical support (ultrafast bootstrap, UFBoot ≥95%) within lineage-specific clades that circulated globally in the post-pandemic period (2023–2025) except one Argentinian strain (VRSV1046), which falls within the pre-pandemic cluster (Fig. 5A and B).

**Fig. 5.**
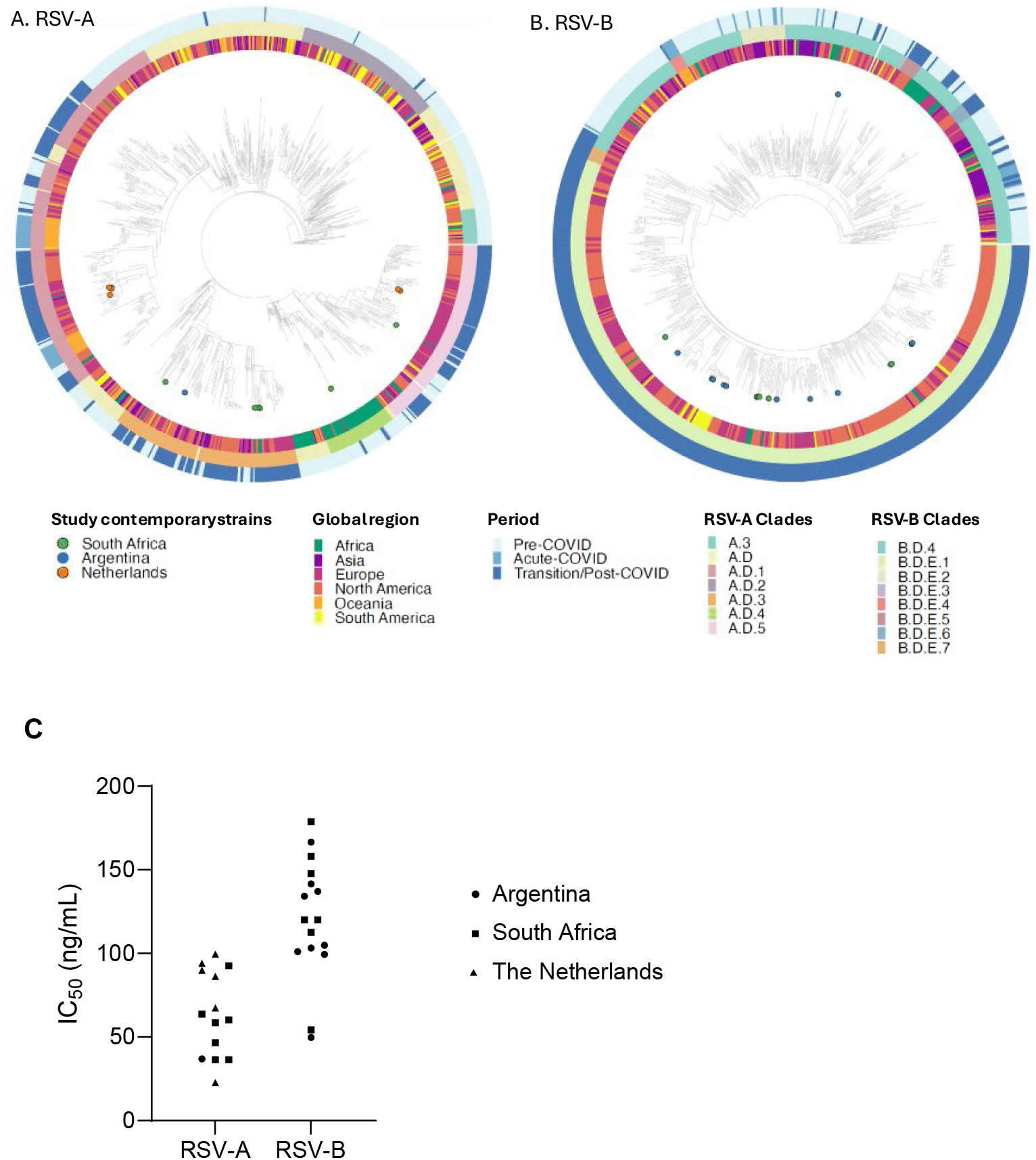
Neutralisation of a diverse panel of global contemporary clinical isolates. The clinical isolates used to evaluate RSM01 neutralisation activity were displayed as a phylogenetic tree to evaluate sequence diversity. **A+B**. Sequences of 14/17 RSV-A and B isolates are marked as coloured dots on clades extracted from a phylogenetic tree of 1,686 RSV-A and 1,650 RSV-B genome sequences representing seven geographic regions representing the time-window from 2010 and 2013, respectively. Pre-pandemic (before 2020), pandemic (2020–2022), and post-pandemic (2023–2025) periods are represented in the outer ring. **C.** RSM01 was assessed for its ability to neutralise RSV clinical isolates using a neutralisation assay. A panel of 30 RSV clinical isolates (14 RSV-A and 16 RSV-B) containing amino acid changes in the F fusion glycoprotein were tested. IC_50_ was calculated and represents the concentration of antibody required for a 50% reduction in the RSV infectivity. IC_50_, half-maximum inhibitory concentration; RSV, respiratory syncytial virus.

RSM01 potently neutralised the RSV-A strains with IC_50_ ranging between 22.7-99.5 ng/mL (geometric mean 58.3 ng/mL) and the RSV-B strains with a range of 49.8-178.8 ng/mL (geometric mean 114.4 ng/mL) (Fig. 5C). Our IC_50_ was reproducible (inter-operator precision 31% coefficient of variation (CV), inter-assay precision 27% CV) and we used an internal control and quality control criteria for every neutralisation assay to assess validity of results.

All of the RSV-B strains carried the S211N mutation without direct association with reduced potency of RSM01 (Table S4). One RSV-B strain carrying an additional R209Q mutation, which is associated with reduced nirsevimab potency, had an IC_50_ of 141.5 ng/mL for RSM01. A South African RSV-A strain carrying two mutations in nirsevimab and palivizumab binding sites (K65R and S276N, respectively) showed no cross-resistance against these mAbs (Table S4). These data confirm that RSM01 is potent against a broad panel of contemporary RSV clinical isolates.

### RSM01 binding to preF

AlphaFold 3 (*26*) predicted that RSM01 binds to the membrane-distal apex of the RSV preF protein, with an interface predicted template modeling score (ipTM) of 0.84, indicative of a high-confidence model (Fig. 6a). The interaction is mediated almost entirely by the CDRH3, which adopts a β-hairpin conformation and forms a three-stranded β-sheet with RSV F residues 210–213 (Fig. 6b). This interaction primarily engages mainchain atoms, and only few contacts with sidechains on RSV F were observed; a notable exception is RSV F lysine 209, which forms two salt bridges with aspartate and glutamate residues in the CDRH3. Observed viral mutations include I206T, which lies underneath the CDRH3, and N216H, which lies outside the direct binding interface but may alter helix capping and influence local dynamics.

**Fig. 6.**
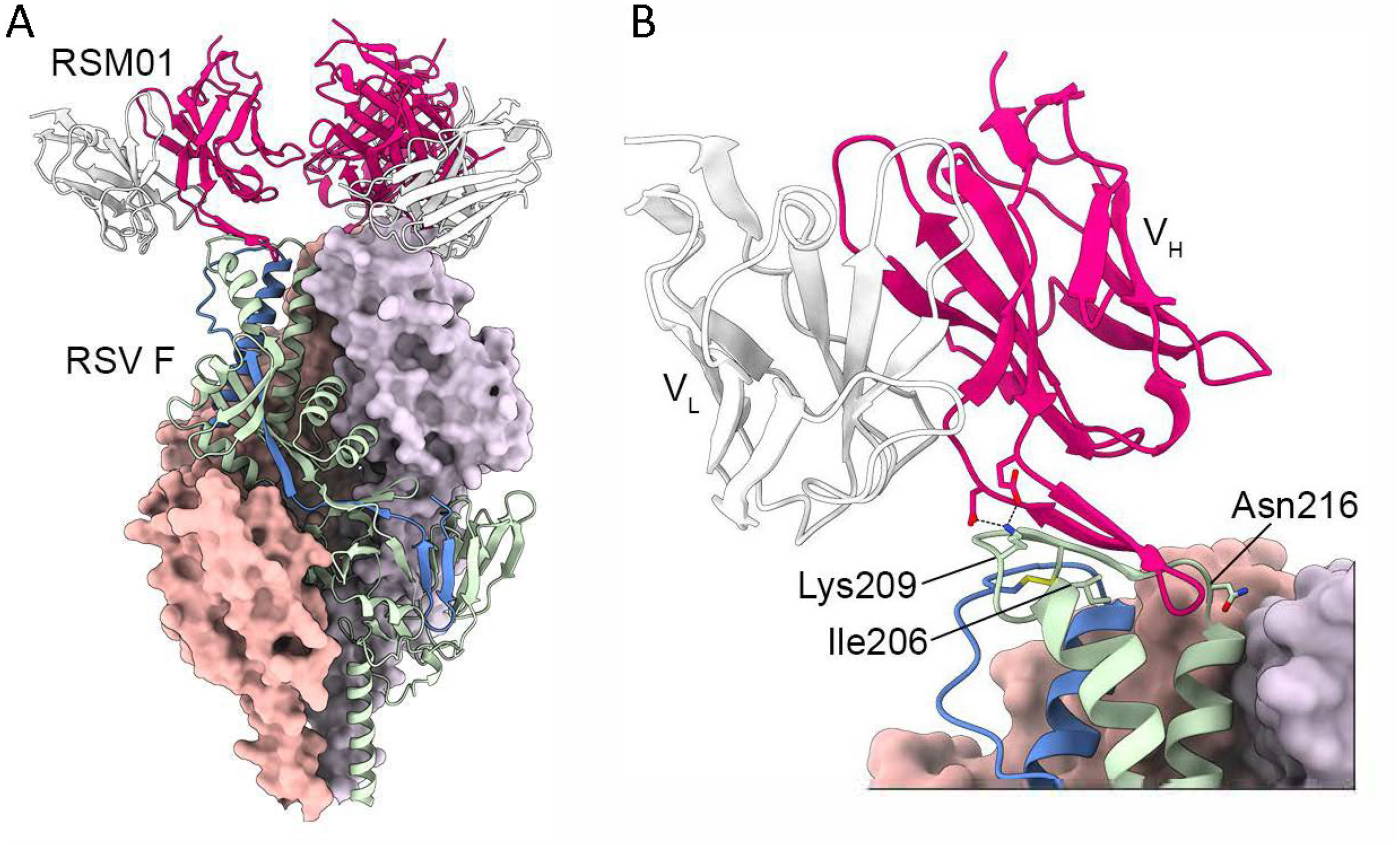
Binding of RSM01 to RSV preF as predicted by AlphaFold. RSM01 binding to the RSV preF protein structure was modeled using AlphaFold. The complex was visualized in UCSF ChimeraX (**A**), highlighting key residues, including viral substitutions at positions 206 and 216 (**B**).

The predicted RSM01 binding mode is similar to that observed for antibody AM22 (*27*), and the targeted region on RSV F corresponds to a metastable segment that contains a conserved disulfide bond (Cys69-Cys212) and undergoes early conformational changes during fusion. The lack of extensive sidechain interactions on RSV F and the metastability of this region may reduce the likelihood of selecting for escape variants that do not have a fitness cost.

## DISCUSSION

In this study, we demonstrate that RSM01 has a high *in vitro* barrier to resistance, confirming previous findings (*8*). Under selective pressure, highly resistant MARMs emerged frequently with palivizumab and nirsevimab, whereas no highly resistant mutants and only low rates of moderate resistance were observed with RSM01. RSM01-selected mutants displayed moderate resistance (10–17-fold IC_50_ increase), likely driven by I206T/N262Y substitutions, compared with markedly higher resistance for nirsevimab- and palivizumab-selected mutants (>1230-fold and >722-fold, respectively). No MARMs showed enhanced growth and one RSM01 mutant exhibited impaired growth. All resistant variants remained susceptible to at least one alternative mAb. A global panel of contemporary strains was susceptible to neutralisation by RSM01.

Previous studies also failed to generate RSM01-resistant mutants, while nirsevimab and palivizumab-MARMs were readily obtained (*8*): whereas palivizumab- and nirsevimab-MARMs emerged within 2– 7 passages, RSM01-MARMs could not be generated in 11 experiments with 8 different RSV strains, suggesting a higher barrier to resistance (Fig. 1C). Contrasting with previous methodology, this study used triple plaque-purified parental RSV strains, viral controls passaged without mAb pressure, systematically generated and identified MARMs, and characterised the MARMs using both functional and sequence data, with virus inoculum that did not violate the principle of mass action. The difficulty in generating RSM01 escape variants suggests that tolerated RSV F substitutions in the epitope are insufficient for escape without substantial fitness costs.

To assess the robustness of RSM01 against globally circulating RSV strains, we assembled a panel of contemporary isolates from 2023–2024 collected across South Africa, Argentina, and the Netherlands. This panel extends previous data (*8*), which included older strains (2002–2017) from a single location (Utrecht, Netherlands). Testing contemporary, geographically diverse isolates is particularly important given potential shifts in RSV strain circulation and antigenicity following the COVID-19 pandemic, which may have altered susceptibility patterns. Preliminary analyses indicate that RSV-B isolates exhibit higher IC_50_s than RSV-A, potentially an artefact of a relatively small sample size or reflecting intrinsic differences in neutralisation sensitivity between subtypes.

We observed higher IC_50_ values for RSM01 against clinical RSV strains (median 99.4 ng/mL) compared with previous work (median 1.8 ng/mL) (*8*), which most likely reflects methodological differences between neutralisation assays, such as multiplicity of infection (MOI) and read-out. Similarly, laboratory RSV-A2 strain against RSM01, nirsevimab, and palivizumab showed an median fold-change in IC_50_ of 16.2 (range 11.8-28.4) between the laboratories. Absolute IC_50_ values vary substantially across laboratories due to differences in reagents, cell density, viral input, overlay, incubation conditions, and readout systems, limiting direct cross-study comparisons. To enable robust interpretation, we therefore assessed resistance of *in vitro* MARMs using fold-changes in neutralisation relative to reference strains within the same experimental setting.

Several mechanisms may explain why RSM01 exhibited a higher barrier to resistance *in vitro*. RSM01 is predicted to engage a highly conserved, backbone-focused epitope at the apex of RSV F, making it largely insensitive to sidechain substitutions that often disrupt other mAbs like nirsevimab. This region is metastable and functionally critical, so mutations that reduce binding likely incur substantial fitness costs by impairing fusion or protein stability. Additionally, RSM01’s functional angle of approach may restrict escape: antibodies requiring multi-residue or conformational changes to evade binding tend to have higher resistance barriers, as seen for RSV, HIV, and influenza (*28*). Together, targeting a conserved, structurally constrained epitope, imposing fitness penalties for mutation, and limiting simple single-point escape pathways likely contribute to RSM01’s observed resistance robustness. Deep mutational scanning could further define which substitutions are tolerated without fitness cost, providing insight into the potential durability of this neutralising antibody.

Some resistance-associated mutations identified in this study have been described previously, but some are novel. For RSM01, we observed a previously described I206T substitution (*29*) within the shared nirsevimab/RSM01 epitope, which functionally affected RSM01 more than nirsevimab. Prior studies reported alternative substitutions at this residue with minimal impact on nirsevimab unless combined with additional changes such as Q209R or N201T (*9, 21, 24, 25*). A novel D194N mutation near the nirsevimab binding site conferred only moderate nirsevimab resistance. We also detected a novel N63K substitution with minimal loss of susceptibility to nirsevimab, similar to other substitutions at this site (N63S clinically; N63T/S *in vitro*) causing minimal IC_50_ increases (*9, 23*). Nirsevimab resistance associated with K65E, reported in both laboratory and clinical strains, disrupts antibody binding via electrostatic repulsion (*23*). Well characterised palivizumab escape mutations N262Y and N272N (> 350-fold increases in IC_50_) were confirmed (*16, 30*). Collectively, these findings show that laboratory-selected MARMs can identify critical preF residues, and alignment with clinical data helps distinguish tolerated mutations from those with potential real-world escape.

These findings have important implications for the global resilience of RSV mAb deployment. In many LMICs, limited genomic surveillance may delay detection of emerging resistance, allowing low-barrier to resistance mAbs to lose effectiveness and worsen health inequities. High-barrier mAbs like RSM01 offer greater resilience, reducing the risk of resistance-driven prophylaxis failure. Moreover, RSM01’s high barrier to resistance provides flexibility for future clinical strategies: it could be combined with other mAbs in bi-specific or cocktail formulations to further limit escape, or potentially used as a salvage prophylaxis in cases resistance renders another mAb ineffective. The specific resistance mutations identified here can also guide targeted surveillance.

This study has several limitations. First, the susceptibility panel included a limited number of contemporary RSV strains, though sourced from three global regions, providing at least partial geographic representation and diversity in a contemporary phylogenetic tree. Second, our genetic analysis was limited by small sample sizes, restricting robust virus population-level statistical comparisons. Non-systematic global surveillance data may have reduced the precision of our assessment of F protein diversity and lineage dynamics over time. Third, plaque-selection of polyclonal selected MARMs potentially caused a loss of MARMs. Preliminary functional data showed no differences in IC_50_ between various plaques of the same polyclonal culture, which led to the practical decision to limit the sequencing of one plaque per MARM. Finally, MARM selection used laboratory-adapted strains rather than recent clinical isolates, which may not capture the full genetic diversity or fitness of circulating viruses and thereby limit the translation of findings to the clinic. However, previous use of clinical strains did not generate MARMs against RMS01 either (*8*). Future studies should validate these findings using a broad panel of contemporary strains in a primary respiratory human cell model to assess resistance in a more physiologically relevant model than immortalized cell lines.

A major strength of this study is the comprehensive panel of RSV mAbs, including licensed and developmental antibodies, enabling direct comparison of resistance profiles. All viral strains were triple plaque purified to ensure genetic homogeneity, and virus controls passaged without mAb showed no mutations, confirming that resistance emerged under selective pressure. A systematic ten-passage protocol, combined with functional data and NGS sequencing, allowed robust identification of resistance-associated mutations, including minor variants. Mass violation studies were performed to assure the neutralisation curves were robust and reproducible. By integrating neutralisation data with global surveillance sequences, we provide functional context to circulating RSV diversity. Detection of multiple co-circulating RSV-A and RSV-B sublineages, alongside their neutralisation phenotypes, improves understanding of temporal patterns and the potential impact of emerging mAb selective pressures.

RSM01 exhibits a high barrier to resistance with rare, moderate, and occasionally fitness-impairing escape mutations. Mapping these resistance pathways highlights key prefusion F residues and informs surveillance strategies. Developing resistance-resilient RSV mAbs is critical to achieving durable, equitable global protection.

## MATERIALS AND METHODS

### Cells, viruses and monoclonal antibodies

HEp-2 cells, RSV-A, and RSV-B were originally obtained from American Type Culture Collection (ATCC). HEp-2 cells (ATCC CCL-23) were passaged in DMEM (Gibco, high glucose, Glutamax, pyruvate, 31966-021) supplemented with 5% fetal bovine serum and pen/strep. Master stocks of RSV-A2 (ATCC VR-1540) and RSV-B1 (B WV/14617/85, VR-1400) were triple plaque purified in HEp-2 cells and propagated for a total of 5 and 8 passages, respectively, before use as parental strains. Four mAbs were evaluated (Table S1): RSM01 (Gates Medical Research Institute), Palivizumab (Synagis®), research-grade Nirsevimab (MedChemExpress, HY-P99756), and research-grade Clesrovimab (MedChemExpress, HY-P99804).

### Generation of monoclonal antibody-resistant mutants

MARMs were generated by serially passaging virus in the presence of palivizumab, nirsevimab, or RSM01. Previous escape mutant selection studies led by AB were performed at Arsanis Biosciences GmbH (Oct 2017 – Nov 2018) according to previously published methodology (*8*). In brief, MARMs were selected by serially passaging laboratory strains and clinical isolates on HEp-2 cells in the presence of fixed concentrations of RSM01 (and variants) or comparator mAbs (nirsevimab (variants) and palivizumab) for a maximum of eight consecutive rounds. Viral populations showing cytopathic effects (CPE) were harvested and F-protein sequences analysed to identify resistance-associated mutations. The growth conditions associated with escape mutations in this earlier work informed the modified selection protocol used in the current study with as main differences that parental strains were triple plaque purified, mAb concentrations were either fixed or rising, and virus controls without mAb pressure.

In the present study performed at the UMC Utrecht, TPP parental strains were mixed with 1-200x published half-maximum inhibitory concentration (IC_50_; Table S1) of mAb and pre-incubated for 30 minutes at 37°C and 5% CO_2_. For simplicity, an IC_50_ of 3 ng/mL was used in nirsevimab concentration calculations during MARM production based on 2 RSV-A lab strains (*8*) and an IC_50_ of 200 ng/mL was used for palivizumab based on lab and clinical strains (*8, 17*). For RSM01, the IC_50_ was set at 2.3 ng/mL based on 2 RSV-A and 2 RSV-B lab strains from previous work by the Gates Medical Research Institute (*8*). HEp-2 cells in 6-well plates were then infected at multiplicity of infection (MOI) of 0.0133-0.00133 plaque-forming units (PFU)/cell. Controls included uninfected cells and virus passaged without antibody pressure. After 1 hour at 37°C and 5% CO_2_, inoculum was removed and fresh medium with appropriate mAb concentrations was added to each well followed by 3-5 day incubation under continuous mAb pressure. The presence of CPE was monitored daily using light microscopy to determine timing of passage, aiming for 40-80% CPE. The contents of each well were harvested by cell scraping and centrifuged at 800g for 5 minutes to remove cell debris. An aliquot of the supernatant was diluted 1:1 with 50% sucrose in dPBS (sterile filter) as a cryo-protectant, snap-frozen, and stored at – 80°C. The remaining supernatant was pre-incubated with fixed or rising concentrations of mAbs and then used to infect HEp-2 cells in the subsequent passage based on the degree of CPE (undiluted virus for no CPE, 1:100–1:1000 virus dilution for low CPE (<50% CPE), and 1:1000 to 1:10.000 virus dilution for high CPE (>50% CPE)). For passaging under rising mAb pressure, virus from the well showing optimal replication (i.e. high CPE) was selected at each passage and subsequently cultured under three mAb concentrations, including the previous and higher concentrations. Concentration increments were empirically adjusted based on viral growth to progressively increase selective pressure. This process was repeated for a total of ten sequential passages. Potential MARMs, i.e. all virus strains cultured for 10 passages under mAb pressure as well as strains showing high CPE under rising mAb pressure at earlier passages, were single plaque purified by infecting HEp-2 cells in 10-fold serial dilutions overlaid with 0.5% agarose to restrict viral spread via supernatant. After 3-5 days, two or three individual plaques were picked and propagated separately under continuous mAb pressure when >10% CPE were visible. After titration, potential MARMs were screened with a neutralisation assay at an MOI of 0.1. If the IC_50_ of a potential MARM was > 3-fold higher than the IC_50_ of the parental strain, it was further characterised for neutralisation capacity after performing mass violation, cross-resistance, RNA mutations, and viral growth assays.

### Tissue culture half infectious dose (TCID_50_) assay

Viral titres were assessed using a tissue culture half infectious dose (TCID_50_) assay as previously described (*31*). In brief, RSV strains underwent quadruplicate 10-fold dilution series in DMEM supplemented with antibiotics and 5% foetal bovine serum for 10 dilutions to infect monolayers of Hep-2 cells. Cells were checked daily for CPE and end-point titres were evaluated as the 50% tissue culture infective dose per mL after 7-10 days using the Spearman-Kärber method (*32*).

### Neutralisation assay

A validated neutralisation assay was adapted to quantify RSV replication by expression of F protein by enzyme-linked immunosorbent (ELISA) assay (*33*). Three-fold serially diluted mAbs were incubated on 96 well-plates with equal volume of RSV for 1 hour at 37°C before adding to a monolayer of HEp-2 cells. After 2 hours, inoculum was replaced by fresh medium with 2.5% FCS and cells were cultured for 3 days at 37°C and 5% CO_2_. Cells were fixed with ice cold 80% acetone in PBS for 15 minutes at 4°C before blocking with 1% BSA in PBS for 1 hour at room temperature. The primary antibody (mouse anti-RSV; Merck Millipore MAB8262) was added 1:2500 for 1 hour and the secondary antibody (goat anti-mouse IgG-HRP; Dako P0447) was added 1:4000 for 1 hour followed by TMB (BioLegend 421101) until colour development was stopped with 1M H_2_SO_4_ after 15 minutes. The absorbance was measured at 450-570 nm (Spectramax M series, Company Molecular Devices). The neutralising titre (IC_50_) was quantified using a log (inhibitor) vs response 4-parameter nonlinear regression curve in GraphPad Prism version 10.6.1 (GraphPad Software Inc., San Diego, CA, USA). An internal control virus neutralised by nirsevimab was taken along on every plate to monitor variation between assays. Other quality control criteria were copied from a previously validated neutralisation assay (*33*).

To assess the law of mass violation (the IC_50_ value obtained in response to a given antibody concentration will remain constant across a range of virus titres (*34*)) and ensure reproducible IC_50_ data, we performed neutralisation assays with RSV strains at a range of three MOIs (*34, 35*). All potential MARMs and contemporary strains were neutralised at an MOI of 2, 0.5 and 0.1, where an MOI of 2 is expected to violate mass action. If an MOI of 2 could not be obtained due to low viral titres, we assessed the highest MOI possible. The principles of mass action were considered violated if the IC_50_ between the highest and lowest MOI differed more than 2-fold. If strains were resistant against one mAb, another mAb was used to determine mass violation. IC_50_ data of the parental strains were obtained by three operators and the MARM IC_50_ data were obtained in replicate by one operator (n=2). IC_50_ data of contemporary strains were obtained by one operator in a single replicate.

Moderate resistance was defined as IC_50_ > 3-fold compared to the parental strain and high resistance > 30-fold IC_50_ compared to the parental strain. Cross-resistance was measured using the neutralization assay against other mAbs.

### Virus growth kinetics

Growth kinetics were performed by inoculating HEp-2 cells with RSV parental strains or MARM variants at an MOI of 0.01 TCID50/cell. The culture supernatant was collected twice daily for four days post-inoculation, and viral titres were determined by a TCID_50_ assay (*31*). Titres below the lower limit of detection (LLOD) were imputed as half the LLOD (158 TCID_50_/mL). Viral growth curves of MARM variants within 1 log deviation of the parental strain were considered as similar growth.

### Contemporary global RSV strains

A panel of 46 newly isolated contemporary global RSV clinical isolates was selected to assess susceptibility to RSM01: 20 strains from South-Africa and 20 from Argentina were prospectively obtained between January and August 2024. Additionally, 6 strains from The Netherlands sampled during Nov-Dec 2023 were randomly selected from the RSV-STRAIN study. Nasopharyngeal swabs were obtained from infants within 48 hours (South-Africa) or 5 days (Argentina and the Netherlands) of hospital admission with PCR-confirmed RSV-associated LRTI and stored at ≤-70°C in universal transport media before shipment on dry ice. If strains were genotyped at time of sample selection, we selected equal amounts of RSV-A and RSV-B strains. Viral stocks were propagated on HEp-2 cells at 37°C and 5% CO_2_ in 1 passage and subsequently sequenced and neutralised by RSM01. One outlier (VRSV1051, RSV-B) with IC_50_ of 337.5 ng/mL was removed from analysis as the neutralisation curve did not meet quality control criteria.

### Sanger sequencing of RSV F-protein

Subtype-specific primer pairs were used to sequence the F-protein. Viral RNA was extracted using Purelink viral RNA/DNA mini kit (Thermo Fisher). 20 μL of RSV samples were isolated according to manufacturer’s protocol and eluted in 50 μL RNase-free water. RT-PCR was performed on 1 μL of RNA sample to produce amplicons of approximately 1000 basepairs from each of the five primer pairs. Thermal cycling protocol was performed on a T100 Thermal Cycler (Bio-Rad) and consisted of a cDNA step at 55°C for 30 min and pre-denaturation at 98°C for 2 min. For amplification 40 cycles of denaturation 98°C for 15 s, annealing 55°C for 15 s, extension 72°C for 1 min, followed by one cycle 72°C for 5 min for final extension. The desired PCR fragments were selected by DNA gel extraction (1% agarose gel). These fragments were further purified by GeneJET Gel Extraction Kit (Thermo Fisher, K0692), according to manufacturer’s protocol, and eluted in 50 μL elution buffer. Samples were sent to Macrogen for Sanger EZ sequencing with sequence primers. Snapgene version 8.0.3. was used to analyse the sequencing data. For data analysis, sequences were aligned with triple-plaque purified parental RSV-A2 and RSV-B1 sequences.

### Next generation whole genome sequencing

#### Nucleic acid amplification

Whole genome sequencing of generated MARMs and global contemporary strains was performed using Illumina NovaSeq X Plus platform, adapted from previously described (*36*). Following RSV subtyping, subtype-specific primer pairs were used to reverse transcribe and amplify four overlapping genome fragments using the SuperScript IV One-Step RT-PCR System (Invitrogen, CA, USA) in a 9800 Fast thermal cycler (Applied Biosystems). These four fragments collectively span the complete RSV genome, encompassing all viral genes. Primer sequences incorporated degenerate bases at positions of genetic variability between RSV-A and RSV-B strains where necessary (*36*). The RT-PCR cycling conditions consisted of reverse transcription at 55 °C for 10 min and initial denaturation at 98 °C for 2 min, followed by 40 cycles of 98 °C for 10 s, 61 °C for 10 s, and 72 °C for 3 min.

Amplicons were verified by electrophoresis on 1% agarose gels, pooled in equimolar ratios, and purified using the GeneJet PCR Purification Kit (Thermo Fisher Scientific). Purified amplicons were quantified using the Quant-iT PicoGreen dsDNA Assay Kit (Thermo Fisher Scientific) according to the manufacturer’s instructions.

#### Library preparation and sequencing

Normalized PCR products were used for library preparation with the Nextera XT DNA Library Prep Kit (Illumina) following the manufacturer’s protocol. Illumina sequencing adapters and unique barcodes were incorporated during PCR amplification of the fragmented DNA using custom oligonucleotide sequences (Integrated DNA Technologies). Tagged DNA was purified and size-selected using 0.6× volume of AMPure XP reagent (Beckman Coulter) according to the manufacturer’s protocol. Purified libraries were quantified with the Quant-iT PicoGreen dsDNA Assay Kit (Thermo Fisher Scientific), pooled in equimolar amounts, and quality assessed using a 4200 TapeStation System (Agilent, USA). Sequencing was performed on either an Illumina NextSeq 500 platform generating paired-end 150 bp reads, or on MiSeq and NextSeq 2000 platforms and whole genome sequence was obtained as previously described (*36*).

#### Viral population level bioinformatic processing and genome assembly

Raw sequencing reads (FASTQ files) underwent quality filtering using Trimmomatic v0.39, with a sliding window approach applied with a 10-base wide window, trimming when average quality dropped below Q30. Leading and trailing bases below Q30 were removed. This process generated paired and unpaired quality-filtered FASTQ files for downstream analysis. The reference sequence was indexed using SAMtools faidx (*37*) for subsequent mapping procedures.

References used for RSV-A and RSV-B respectively, Sequences EPI_ISL_412866 (hRSV/A/England/397/2017) and EPI_ISL_1653999 (hRSV/B/Australia/VIC-RCH056/2019). Quality-filtered paired-end reads were mapped to the reference genome using Minimap2 v2.26 with the short-read alignment mode. Variant calling was performed using LoFreq (*38*). Minor variants were filtered to retain only those with allele frequency (AF) 0.4<AF≥0.05 and sequencing depth (DP) ≥200 using BCFtools filter (*37*) and customized scripts with minMutFinder (*39*). Variant annotation in variant call format (VCF) was performed using SnpEff with a custom-built database based on the reference genome (*40*) and GFF3 annotation file. Samples were aligned to reference sequences using Nextclade align, translated into amino acids, and annotated for mutation profiles using Nextclade v3.18.1 (*41*). To identify polymorphisms in the F protein, binding regions targeted by RSV mAbs were then screened for mutations. MARMs were named after the mAb used to generate the MARM and numbered (RSM for RSM01, NIR for nirsevimab, and PAL for palivizumab).

### Prevalence of resistance-associated mutations and Shannon entropy

To assess the prevalence of observed resistance-associated mutations in circulating strains as well as genetic diversity within RSV-A and RSV-B F-protein sequences at mAb binding sites, RSV genomes deposited in GISAID between May 5, 2023, and December 1, 2025 (post-COVID period) were retrieved on December 12, 2025. Genomic sequences were aligned using Nextclade align and annotated using Nextclade v3.18.1 (*41*), yielding 5,037 RSV-A and 5,120 RSV-B F-gene sequences.

Shannon entropy (H) was calculated to quantify amino acid diversity at each position within the F protein (*42*). Reference strains were excluded from the analysis, and aligned sequences were converted to a character matrix. The frequency of each amino acid at each position was determined, and Shannon entropy (H) was computed according to the following formula:

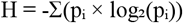

where p_i_ represents the frequency of amino acid i at a given position. Higher entropy values (H > 0.1) indicate greater amino acid variability (*43*) while lower values reflect conservation. Shannon entropy estimates for RSV-A and RSV-B were visualised using the Matplotlib library (Python).

### Phylogenetic reconstruction

To reconstruct phylogenetic relationships among contemporary RSV clinical strains and assess the temporal dynamics of circulating lineages, we obtained complete, high-quality RSV genome sequences spanning 1950 to 2025 from the Nextstrain database (*44*) via Pathoplexus. Sequences were annotated using Nextclade v3.18.1 (*41*), yielding a dataset encompassing diverse genomic regions and clades. The final dataset comprised 1,686 RSV-A and 1,650 RSV-B genome sequences representing the seven geographic regions and was structured to capture temporal patterns across three epidemiological periods: pre-pandemic (before 2020), pandemic (2020–2022), and post-pandemic (2023–2025). The maximum-likelihood phylogenetic inference was performed with IQ-TREE v3.01 and branch support was assessed using an ultrafast bootstrap approximation (UFBoot with 1,000 replicates). Final phylogenetic trees with bootstrap support values were visualised using the ggtree R package (*45*).

### Structural modelling of RSM01-preF complex

The 3D structure of the RSM01 variable heavy and light chains in complex with the RSV F ectodomain was predicted from amino acid sequences in a 3:3:3 ratio using the AlphaFold 3 server accessed on January 23, 2026 (*46*) with default parameters. The highest-confidence model was selected based on interface predicted TM (ipTM)-scores and predicted aligned error metrics. Structural models were visualized using UCSF ChimeraX (version 1.10) (*47*).

## Ethical approval

The study protocols were approved by the Institutional Review Board at Hospital General de Niños Pedro de Elizalde, Argentina (approval number 12863) and the Human Research Ethics Committee at the University of Witwatersrand, South Africa (approval number M170966). The institutional research office of the University Medical Centre Utrecht (Object ID 23U-0590) provided a waiver for the STRAIN study as the Dutch Medical Research Involving Human Subjects Act was not applicable. Written informed consent was obtained from parents or guardians of patients before inclusion in the study.

## LIST OF SUPPLEMENTARY MATERIALS

Table S1-4

Fig. S1-4

Additional references (*48-50*)

## NOTES

## Acknowledgments

We thank Anouk Evers for performing all the deep sequencing and Barney S. Graham for his valuable advice. ChatGPT was used as a language editor to optimise the flow of the manuscript.

## Author contributions

Conceptualisation: JT, LJB, AB, NIM

Data curation: JT, EGB, RJL

Formal analysis: JT, EGB

Funding acquisition: JT, LJB

Investigation: JT, JSM, KC, JD, RJL, ICEL, JR, AV, MV

Methodology: JT, EGB, JSM, KC, RJL, AV, EMD, AB, NIM

Project administration: EMD, NIM

Resources: VLB, MTC, JD, JR, MV, HJAAZ

Supervision: LJB, EMD, NIM

Visualisation: JT, EGB, JSM, ICEL

Writing – original draft: JT, EGB, JSM, NIM

Writing – Review & Editing: all co-authors

## Data and materials availability

The genomic sequences generated during this study are available in the open-source Pathoplexus database (accession numbers **RSV-A**: PP_004HWQJ.1, PP_004HWRG.1, PP_004HWSE.1, PP_004HWTC.1, PP_004HWUA.1, PP_004HWV8.1, PP_004HWW6.1 and PP_004HWX4.1; **RSV-B**: PP_004HXG0.1, PP_004HXHX.1, PP_004HXJV.1, PP_004HXKT.1, PP_004HXLR.1, PP_004HXMP.1, PP_004HXNM.1, PP_004HXPK.1, PP_004HXQH.1, PP_004HXRF.1, PP_004HXSD.1, PP_004HXTB.1, PP_004HXU9.1, PP_004HXV7.1, PP_004HXW5.1, PP_004HXX3.1, PP_004HXY1.1, PP_004HXZZ.1).

Data sources files and scripts developed for this study are available in a dedicated Github repository: github.com/emanuelegustani/Next-Generation_RSV_Monoclonal-Antibody_with_high_barrier_to_resistance.git

