## Supplementary for "RSM01, an extended half-life RSV monoclonal antibody with a high barrier to resistance"

This appendix has been provided by the authors to give readers additional information about the work.

### Supplementary figures

**Fig. S1.** RSV monoclonal antibodies and their binding sites on RSV preF

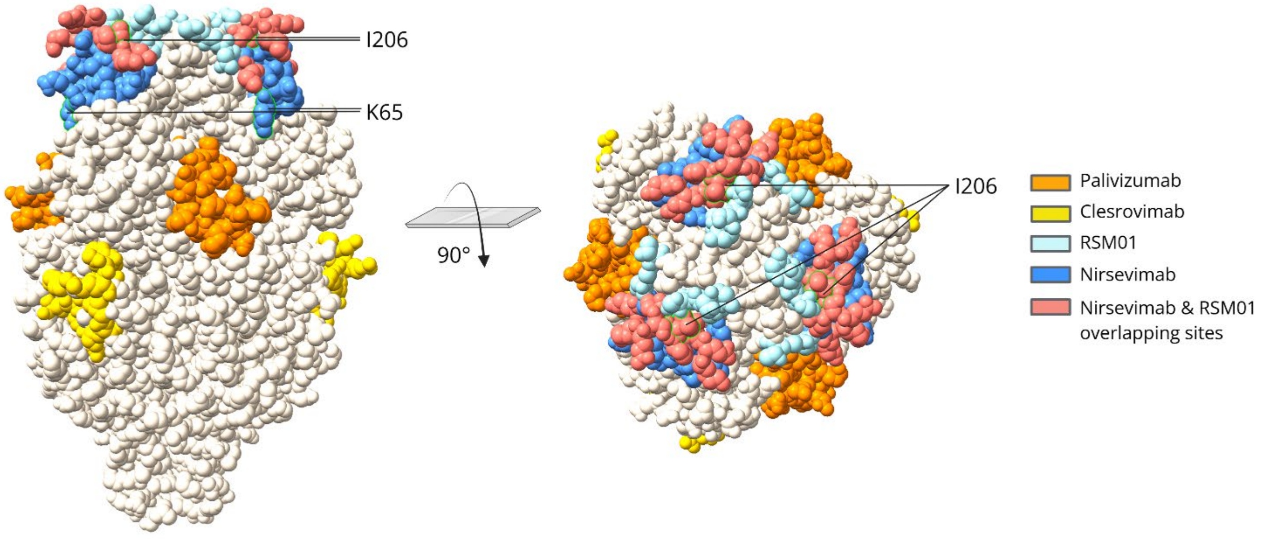


RSV Fusion protein in prefusion confirmation. Palivizumab (orange) targets antigenic site II, while clesrovimab (yellow) targets antigenic site IV. RSM01 (light blue) and nirsevimab (dark blue) epitopes overlap (salmon) in antigenic site Ø. Figure was produced in ChimeraX-1.10.1 using RCSB Protein Date Bank 8YE3 (*48, 49*). PreF, prefusion protein; RSV, respiratory syncytial virus.

**Table S1.** RSV monoclonal antibody characteristics

| **mAb** | **Antigenic site** | **preF epitopes (amino acid position)** | **Published $IC_{50}$ (ng/mL)** |
|---|---|---|---|
| RSM01 | ø | 68, 70-72, 201-203, 205-206, 209-216, 219 (*8*) | 1.7 (RSV-A) and 2.2 (RSV-B) (*8*) |
| Nirsevimab | ø | 62–69 and 196–212 (*17, 21*) | 2.1 (RSV-A) and 2.4 (RSV-B) (*17*) |
| Palivizumab | II | 262-275 (*50*) | 650 (RSV-A) and 280 (RSV-B) (*50*) |
| Clesrovimab | IV | 426-429, 432-433, 440-441, 443, 445-447 (*18*) | 3.7 (RSV-A) and 4.5 (RSV-B) (*18*) |

Antigenic site and binding epitopes are on RSV prefusion (preF) protein. The $IC_{50}$ is estimated based on available literature taking the median $IC_{50}$ of a mixture of laboratory and clinical strains. $IC_{50}$, half-maximum inhibitory concentration; mAb, monoclonal antibody; MARMs, monoclonal antibody resistant mutants; RSV, respiratory syncytial virus.

**Fig. S2.** Flow chart of monoclonal antibody resistant mutant generation

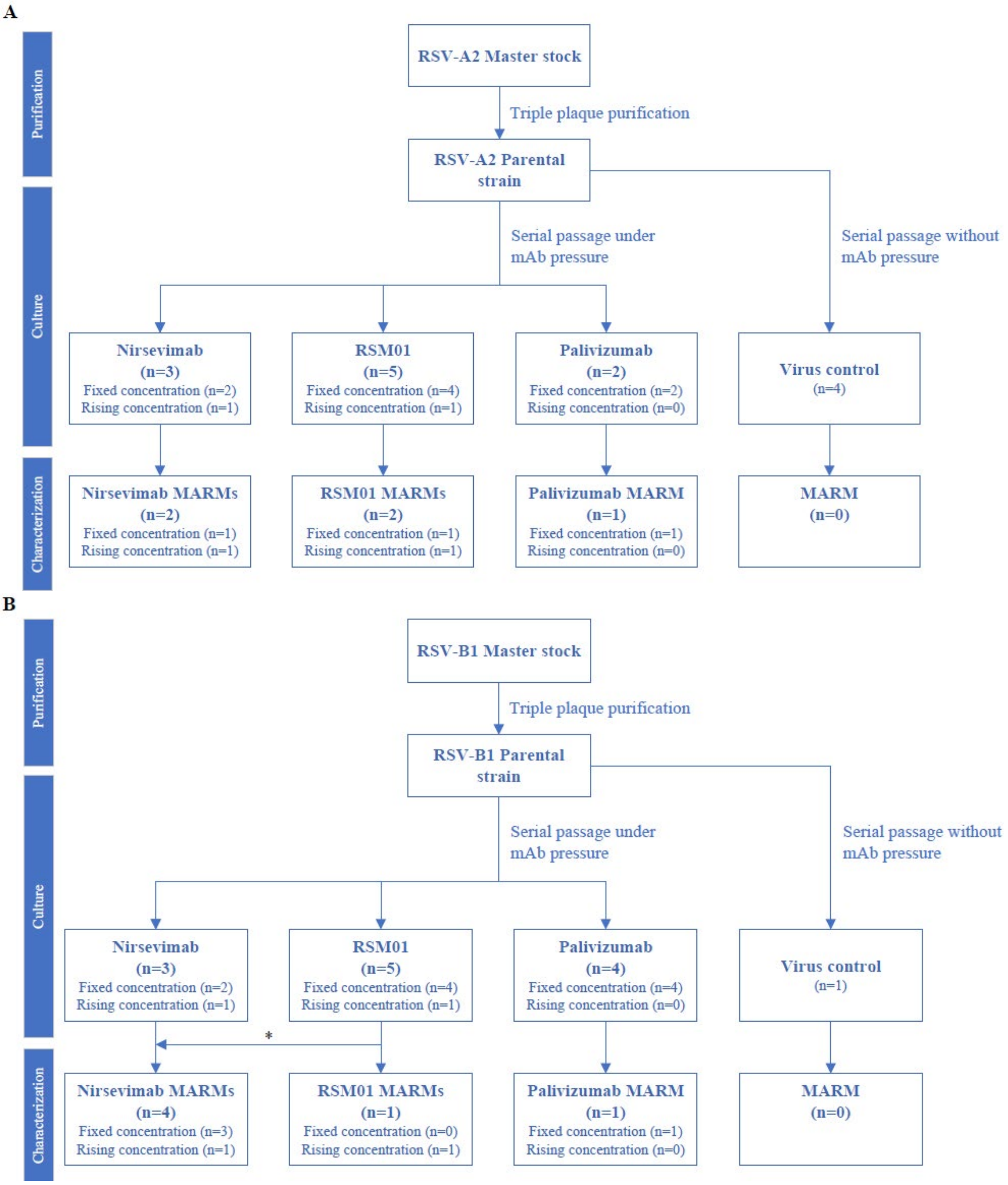


RSV-A2 (**A**) and RSV-B1 (**B**) ATCC master stocks were triple plaque purified to obtain the parental strains. Monoclonal antibody resistant mutants (MARMs) were generated by serial passage of parental virus in the presence of mAbs (palivizumab, nirsevimab, or RSM01). The concentration of mAb was either fixed for 10 passages or rising based on the degree of CPE observed via light microscopy. As a control, the parental strains were also serially passaged without mAb pressure to ascertain that resistance occurs due to mAb presence rather than serial passage. (**A**) RSV-A2 was cultured under 10 mAb conditions, which generated 5 MARMs. (**B**) RSV-B1 was cultured under 12 mAb conditions, which generated 6 MARMs. *One of the RSM01 cultured MARMs, showed a mutation in the nirsevimab epitope and was also characterized into a highly resistance mutant The virus controls showed no mutations compared to the parental strain. CPE, cytopathic effects; mAb, monoclonal antibody; MARM, monoclonal antibody resistance mutant; RSV, respiratory syncytial virus.

**Figure S3.** Viral growth kinetics of master stocks and the triple plaque purified parental strains on HEp-2 cells

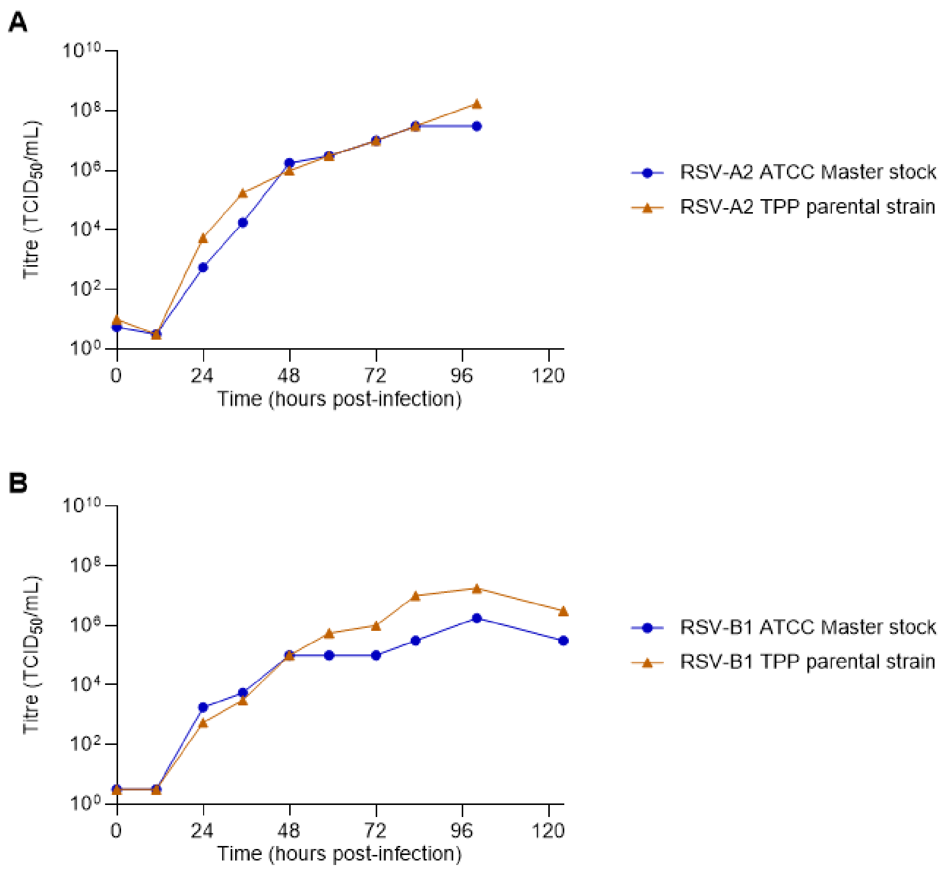


Growth kinetics of ATCC master stocks RSV and the derived triple-plaque purified (TPP) parental strains. HEp-2 cells were inoculated at an MOI of 0.01 $TCID_{50}$/cell. The cultured supernatant was sampled twice daily for five days and viral growth was assessed using a $TCID_{50}$ assay ($TCID_{50}$/mL). (**A**) RSV-A2 master stock (ATCC+2 passages) and TPP-parental strain (ATCC+7) and (**B**) RSV-B1 master stock (ATCC+1) and TPP-parental strain (ATCC+9) show similar growth kinetics.

**Table S2.** Amino acids positions of the fusion protein gene with high entropy (H > 0.1) among RSV-A isolates.

| AA Position | Entropy |
|---|---|
| 12 | 0.94 |
| 122 | 0.84 |
| 377 | 0.65 |
| 127 | 0.58 |
| 103 | 0.52 |
| 276 | 0.51 |
| 23 | 0.38 |
| 20 | 0.30 |
| 3 | 0.28 |
| 15 | 0.27 |
| 114 | 0.23 |
| 518 | 0.21 |
| 22 | 0.20 |
| 123 | 0.20 |
| 543 | 0.18 |
| 119 | 0.17 |
| 105 | 0.17 |
| 17 | 0.16 |
| 384 | 0.14 |
| 523 | 0.13 |
| 115 | 0.13 |
| 107 | 0.12 |
| 63 | 0.11 |
| 120 | 0.11 |
| 470 | 0.11 |

Entropy was defined as $-\Sigma(p_i \times \log_2(p_i))$ and amino acid (AA) positions were ranked by entropy value.

**Table S3.** Amino acids positions of the fusion protein gene with high entropy ($H > 0.1$) among RSV-B isolates.

| AA Position | Entropy |
|---|---|
| 42 | 0.76 |
| 389 | 0.75 |
| 12 | 0.56 |
| 190 | 0.40 |
| 209 | 0.36 |
| 529 | 0.31 |
| 103 | 0.26 |
| 211 | 0.25 |
| 327 | 0.24 |
| 115 | 0.23 |
| 19 | 0.21 |
| 13 | 0.18 |
| 118 | 0.17 |
| 125 | 0.15 |
| 517 | 0.15 |
| 462 | 0.11 |
| 6 | 0.11 |

Entropy was defined as $-\Sigma(p_i \times \log_2(p_i))$ and amino acid (AA) positions were ranked by entropy value.

**Table S4.** Functional and genetic characterisation of global contemporary strains

| Country of origin | Strain | RSV subtype | Clade | $IC_{50}$ (ng/mL) | F-protein mutation(s) |
|---|---|---|---|---|---|
| SA | PET31333 | A | A.D.5.1 | 36.5 | A23S, T122A |
| | PET31973 | A | A.D.3 | 46.5 | T12I |
| | PET31976 | A | A.D.3.1 | 36.4 (R)<br>117.8 (N)<br>7372 (P) | T12I, L20F, K65R, S276N |
| | PET32157 | A | A.D.3 | 60.3 | A10T, T12I, S25C |
| | PET32437 | A | A.D.3 | 92.6 | T12I |
| | PET33105 | A | A.D.3 | 58.5 | T12I, K390E, L547F |
| | PET33037 | A | A.D.4.1 | 63.7 | T122A, K123Q, S377N, I384T |
| | PET32029 | B | B.D.E.1 | 112.7 | A19T, S190N, S211N, S389P |
| | PET32021 | B | B.D.E.1 | 54.2 | R42K, S190N, S211N, S389P |
| | PET32254 | B | B.D.E.1 | 120.1 | R42K, S190N, S211N, S389P |
| | PET32338 | B | B.D.E.1 | 178.8 | A19T, S190N, S211N, S389P |
| | PET31620 | B | B.D.E.1 | 120.1 | S190N, S211N, S389P |
| | PET31686 | B | B.D.E.1 | 147.8 | S190N, S211N, S389P |
| | PET32766 | B | B.D.E.1 | 158.0 | A19T, S190N, S211N, S389P |
| AR | VRSV1056 | B | B.D.E.1 | 101.1 | A107V, S190N, S211N, S389P |
| | VRSV1042 | B | B.D.E.1.1 | 134.2 | S190N, S211N, S389P |
| | VRSV1041 | A | A.D.3.3 | 36.9 | T12I |
| | VRSV1036 | B | B.D.E.1.7 | 166.6 | R42K, S190N, S211N, S389P |
| | VRSV1050 | B | B.D.E.1 | 103.3 | S190N, S211N, S389P |
| | VRSV1048 | B | B.D.E.1.1 | 99.4 | T122I, S190N, S211N, S389P |
| | VRSV1045 | B | B.D.E.1.1 | 49.8 | T122I, S190N, S211N, S389P |
| | VRSV1035 | B | B.D.E.1 | 141.5 | S190N, R209Q, S211N, S389P, V406I |
| | VRSV1031 | B | B.D.E.1 | 104.9 | L22P, S190N, S211N, S389P |
| | VRSV1043 | B | B.D.E.1.1 | 137.1 | S190N, S211N, S389P |
| | VRSV1051 | B | B.D.E.1.7 | 337.5 | R42K, S190N, S211N, S389P |
| NL | RSV-STRAIN 03 | A | A.D.1.9 | 67.4 | S377N |
| | RSV-STRAIN 05 | A | A.D.1.9 | 22.7 | S377N |
| | RSV-STRAIN 13 | A | A.D.5.2 | 89.8 | A103T, M115T, T122A, R553K |
| | RSV-STRAIN 20 | A | A.D.1.9 | 99.5 | S377N |
| | RSV-STRAIN 23 | A | A.D.1.9 | 94.1 | S377N |
| | RSV-STRAIN 27 | A | A.D.5.2 | 86.2 | A103T, M115T, T122A, R553K |

A panel of 31 RSV clinical isolates (14 RSV-A and 17 RSV-B) containing amino acid changes in the F fusion glycoprotein were tested in the neutralisation assay. $IC_{50}$ was calculated and represents the concentration of antibody required for a 50% reduction in the RSV infectivity. Presented $IC_{50}$ for RSM01 unless stated otherwise (where N annotates nirsevimab, P palivizumab, and R RSM01). F-protein mutations are relative to the reference strain (see supplementary methods). Abbreviations: AR, Argentina; $IC_{50}$, half-maximum inhibitory concentration; NL, the Netherlands; RSV, respiratory syncytial virus; SA, South-Africa

**Fig. S4.** Diversity of global contemporary clinical isolates against globally representative sequences from six geographic regions spanning 1950–2025

**A.** RSV-A

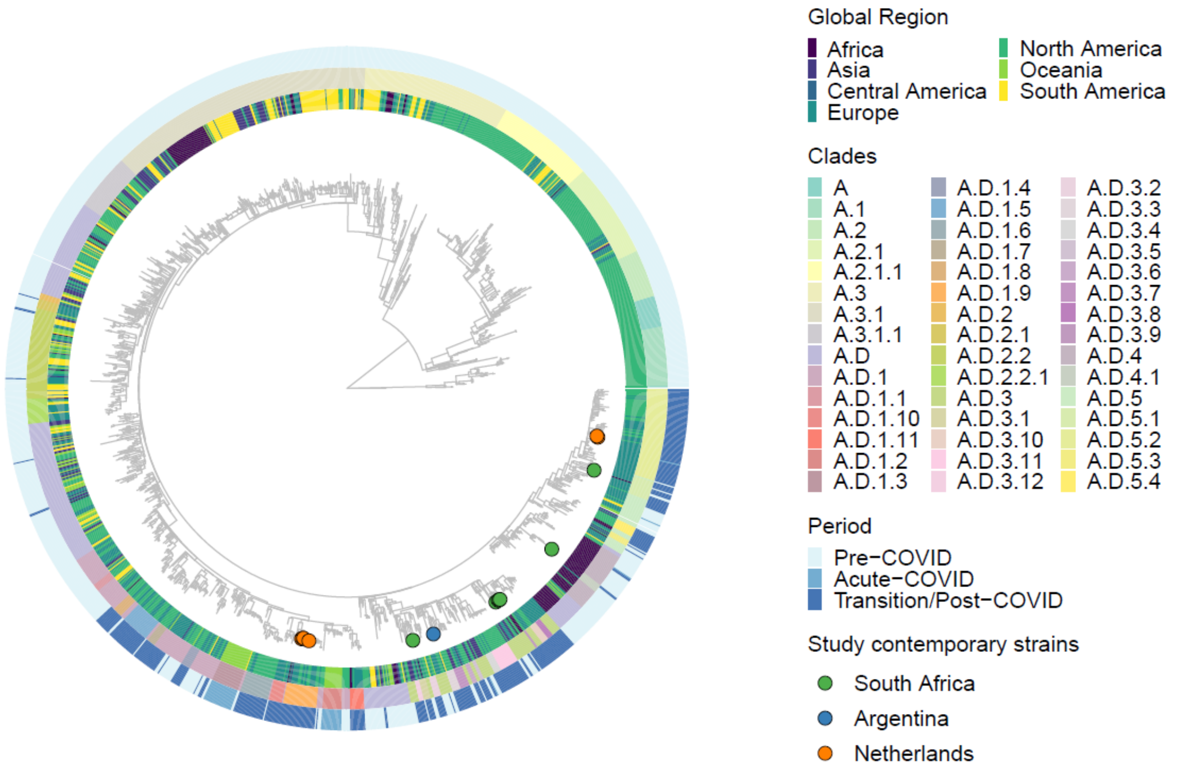


**B.** RSV−B

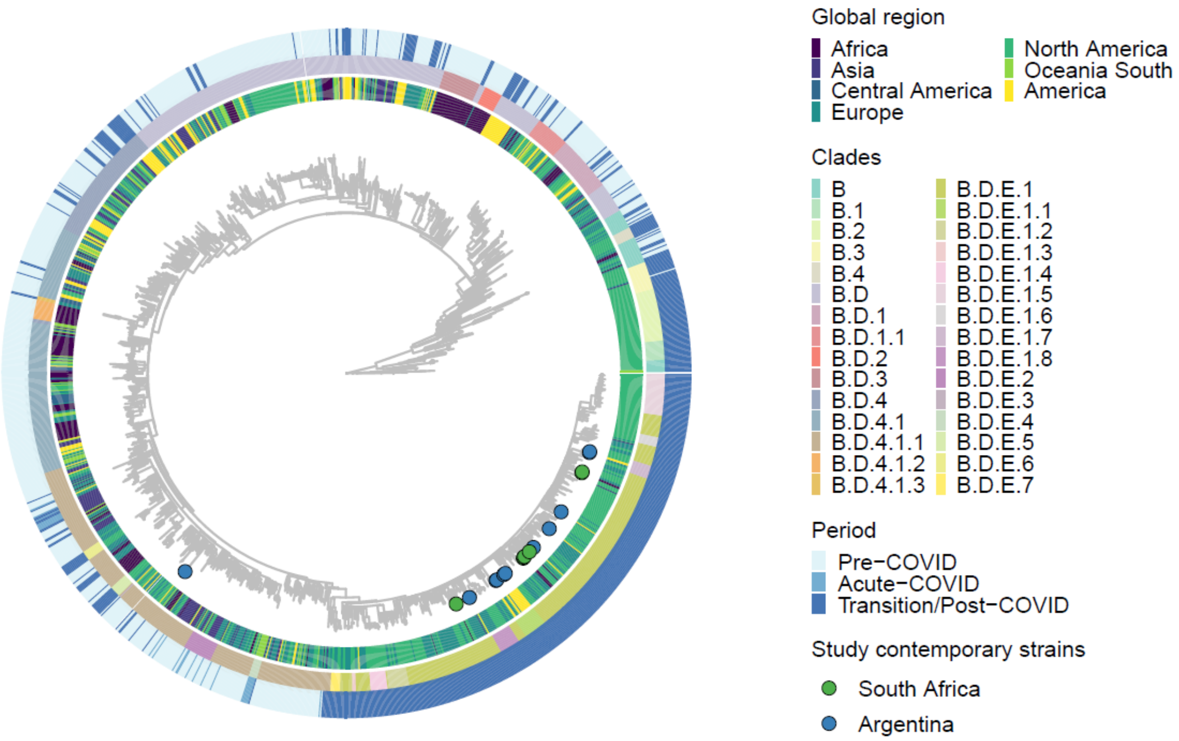


Phylogenetic tree of 1,686 RSV-A (**A**) and 1,650 RSV-B (**B**) genome sequences representing seven geographic regions representing the time-window from 1950 to 2025. Sequences of 14/17 RSV-A and B isolates are marked as coloured dots. Pre-pandemic (before 2020), pandemic (2020–2022), and post-pandemic (2023–2025) periods are represented in the outer ring.
